# Becoming Biomedical Faculty: A Longitudinal Analysis of Successful Academic Career Aspirants’ Career Perspectives, Motivations, and Intentions

**DOI:** 10.64898/2026.05.20.726590

**Authors:** Remi F. Jones, Cilka M. Hijara, Christine V. Wood, Robin Remich, Patricia B. Campbell, Anne E. Skelley, Julia F. Mendes, Yun Kyung Cho, David P. O’Neill, Richard McGee

**Affiliations:** Feinberg School of Medicine, Northwestern University, Chicago, IL, USA; QEM Network, Washington D.C., USA; Campbell-Kibler Associates, Inc., Groton, MA, USA; Northwestern University, Chicago, IL, USA; Robert R. McCormick School of Engineering and Applied Science, Department of Biomedical Engineering, Northwestern University, Evanston, IL, USA; Feinberg School of Medicine Faculty Affairs, Northwestern University, Chicago, IL, USA

**Author notes:** These authors contributed equally to this work (Remi F. Jones, Cilka M. Hijara).

## Abstract

Seismic shifts within academia over the last several decades have seen the growth of biomedical PhD recipients alongside the relative stagnation of tenure-track research-intensive faculty careers (RIFCs). This hypercompetitive academic job market has prompted interest in the paths of those who attain RIFCs. Understanding what drives recent biomedical PhDs to make their career decisions and persist toward them requires a clear picture of how career perceptions, motivations, and intentions develop and crystallize over time. Using annual in-depth interviews across nearly two decades, this report explores the evolution of career thinking and differentiation among 40 who attained a RIFC from diverse starting points to their attainment of a RIFC. Participants’ strategies for navigating early scientific experiences were patterned by their varied educational and socioeconomic backgrounds. Nearly half of participants did not start with or maintain stable interest in RIFCs, exhibiting changes in both PhD and postdoctoral phases. Participants highlighted six ‘drivers’ toward RIFCs including desire for independence/autonomy and contributing to knowledge/health. Our results are instructive for trainees and mentors guiding career exploration and differentiation.

## 1. Introduction

As of 2024, there were 62,357 biomedical PhD students at various stages of training in U.S. programs (1). Over the years spent in training, these tens of thousands evolve and differentiate into a wide array of careers using skills acquired during training. Both internal (e.g., scientific development, motivations) and external (e.g., mentorship, external validation) forces inform career differentiation. Yet the factors behind this process of differentiation have never been studied in detail on a longitudinal basis. Of the many who enroll, approximately 9,000-10,000 complete their doctoral training in biomedical sciences annually, going on to seek postdoctoral fellowships or employment (2). In recent decades, the faculty job landscape has become increasingly competitive with the growing number of postdoctoral positions and PhDs vastly outpacing the available tenure-track positions (3). These seismic shifts toward careers outside of academia have brought increased interest in those who attain tenure-track research-intensive faculty careers (RIFCs). Thus, it is critical to understand what drives recent biomedical PhDs to make their career decisions.

Work that explores the trajectory of interest and intention has revealed that biomedical PhDs frequently lose interest in becoming PIs over the course of PhD training (4–6). However, the decline of interest in PI positions is not universal. Studies also report a subset whose intention towards a PI career increase as well as those who have a steady intention towards a PI career from the start of PhD training (5, 6). Moreover, few qualitative studies have attempted to explain the dynamic, multi-factorial process behind RIFC career differentiation and persistence, largely examining only one factor at a time.

Our research team has maintained a national longitudinal study of biomedical research trainees for close to twenty years, tracking participants from the transition out of undergraduate education / the start of postbaccalaureate or graduate programs through the early career period, using annual in-depth interviews. These individuals started graduate training between 2006 and 2011. Data collection is still ongoing, with the majority of participants now navigating early career positions and a small number still completing postdoctoral training. Our previous analyses looked closely at the formation of and changes in career intentions over the course of PhD training, finding that the career decision-making process is highly fluid and most biomedical graduate students change their career goals several times before graduation (6). Our current analysis of longitudinal interviews reveals more about the career differentiation process by extending the timescale to include postdoctoral years and career attainment.

Additionally, we were able to examine whether and to what extent career intentions continue to be fluid during postdoc positions. We undertook thematic analysis that revealed varied processes that shape career differentiation throughout both PhD and postdoctoral years. These findings can help trainees assess their motivations, values, and perspectives, helping prepare them for career decision. With an increased understanding of career intention patterns and what fuels persistence, mentors can better support their trainees along varied paths towards RIFCs and other careers. RIFCs are the most commonly cited goal upon starting a PhD, so understanding how those who attain them progressed and differentiated into those careers is a vital part of the career differentiation puzzle.

Using a sample of 40 trainees who participated in a total of 393 interviews (see Methods for discussion of sample size), this report examines the processes that shaped differentiation into RIFCs. With individuals interviewed over many years, we can ask the following questions: What factors and pathways lead to the decision to pursue this career, and what variability exists across individuals? What patterns of intention exist? What motivations do those who attain a RIFC hold highest? What extrinsic and intrinsic factors do individuals cite in their decision to pursue a RIFC and how do these factors interact? What promotes individual persistence towards a highly rarefied career?

## 2. Materials and Methods

### Description of the Population and Data Sources

This study’s population came from two different NIH-funded longitudinal studies, the National Longitudinal Study of Young Life Scientists (NLSYLS) and the Academy for Future Scientists (the Academy), that investigate scientific and professional development alongside career differentiation of a national population of biomedical graduate students. Researchers conducted annual, one-on-one interviews and surveys. Northwestern University’s institutional review board approved both studies.

Seventeen of the 40 participants in this subsample were enrolled in the NLSYLS. The NLSYLS is a longitudinal, qualitative, interview-based study of biomedical research trainees’ career differentiation, following from the start of their PhD programs through early-career stages. From 2008 to 2011, we recruited participants from colleges and universities across the US and Puerto Rico. Different waves of recruitment included a cohorts of late-stage undergraduates and postbaccalaureate students intending to enter graduate school as well as beginning PhD students. The NLSYLS annually interviewed 260 biomedical PhD students, discontinuing those who chose non-academic careers, those who left their PhD programs, and those who voluntarily withdrew. Two-hundred-and-eight of the 260 graduated from their PhD programs. Twenty-eight of the 260 are currently in postdoctoral training. A forthcoming publication will display the career outcomes of the 208 who completed the PhD.

In this subsample, 23 of 40 participants are participants from “the Academy.” Initiated in 2011, the Academy began as theory-based, career-coaching intervention focused on persistence towards RIFCs. The Academy consisted of two cohorts: Academy 1’s population consisted of students nearing the start of their PhD programs, and Academy 2’s population consisted of participants nearing PhD defense and graduation. “Coaches” were recruited from the ranks of senior biomedical faculty who possessed exemplary competencies in research training, mentoring, and career development served. We annually interviewed Academy participants covering experiences related to their time in the Academy as well as graduate, postdoctoral, career search and attainment phases.

Both Academy 1 (n=180) and Academy 2(n=117) were randomized into two arms, an experimental arm who received the coaching intervention, and a control arm without coaching. Interventions entailed three years of both in-person and virtual coaching sessions. This report draws only on interview data; the results of the Academy intervention have been reported previously (27).

We were able to merge the NLSYLS and the Academy with support from an R35 MIRA award. The interview questions covering training and career differentiating were nearly identical in both studies; we consequently can treat data from both studies as comparable. Interview guides have been made available in an earlier publication (28).

The original population of those who attained RIFCs included 54 participants. Fourteen did not complete an adequate number of interviews to make detailed analysis possible, and they are consequently not included in this report.

Table 1 displays this subsample’s sociodemographic characteristics. While there are an identical number of men and women, the total population in our study consists of a higher proportion of women (61%). Similarly, Black/African American and Hispanic individuals each represent 12.5% of the 40. This number is a higher proportion than new biomedical faculty hires (7). In our entire study population, 17% of doctoral recipients are Black/African American and 18% are Hispanic much higher than the respective 3% and 9% reported by the NSF (2). Thirteen reported growing up with less than the mean household family income, while fifteen reported growing up in one of the two highest income categories. Parents’ highest degree of education was likewise varied, a quarter had parents with a high school education or less, another quarter held a bachelor’s degree. Nine were either first-generation immigrants (n=4) or the children of immigrants (n=5). Our sample of individuals who attained a RIFC includes a variety of backgrounds.

**Table 1.** Participant Sociodemographic Characteristics.

|  |  |
| --- | --- |
| Gender |  |
| Women | 20 |
| Men | 20 |
| Race/Ethnicity |  |
| White | 25 |
| Black/African American | 5 |
| Hispanic, Latino/a* | 5 |
| Asian | 4 |
| Multiple | 1 |
| Immigration Generation |  |
| First-Generation | 4 |
| Child of Immigrants | 5 |
| Parents' highest degrees |  |
| High School/GED or less | 10 |
| Baccalaureate degree | 10 |
| MS/PhD/Professional | 20 |
| Family income in 10<sup>th</sup> grade |  |
| Under \$20,000 | 3 |
| \$20,000-\$44,999 | 5 |
| \$45,000-\$64,999 | 5 |
| \$65,000-\$89,999 | 9 |
| \$90,000-\$124,999 | 7 |
| \$125,000 plus | 8 |
| unknown | 3 |
| Median family income in 2000<sup>†</sup> | \$51,710 |
| Mean family income in 2000<sup>†</sup> | \$66,710 |

\*Original survey text asked participants if they identify as non-White Hispanic/Latino.
<sup>†</sup>We include median and mean family income in 2000 (the approximate year that many participants would have been in 10<sup>th</sup> grade) to contextualize participants' reported family income.

### Data Collection and Analysis

The initial and second NLSYLS annual interviews were in-person, taking place at each participant’s university. Following interviews were conducted via telephone and later by Zoom. All Academy interviews were conducted via telephone or Zoom. For both Academy and NLSYLS, interview protocols covered topics of pre-PhD science and research exposure; PhD program, lab, and dissertation selection; PhD and postdoctoral research experiences; choices of mentors and subsequent experiences of mentoring; career perspectives and intentions as well as other future plans. This annual interview methodology was able to gather real-time data at regular intervals. As of the writing this manuscript, the median number of interviews was 10.

We utilized a thematic analysis strategy to arrive at saturation, a well-supported metric for assessing completeness of analysis and adequacy of sample size in qualitative research. We analyzed 393 interviews across the population of 40 participants. Thematic saturation is arrived at when no additional themes are uncovered, and researchers are empirically confident that conclusions are thoroughly developed (8–10).

Much smaller sample sizes than we have (e.g., 9-12 interviews) are typically sufficient to reach s (10–11). In our population, we approached saturation—identifying themes, their meanings, importance, and their relationships—after analysis of approximately 40 interviews of10 participants. To increase our empirical confidence and to base conclusions on a full range of participants from various demographic backgrounds, we continued analysis beyond saturation.

The research team created an NVivo database where full transcripts were coded. Initial coding utilized broad descriptive and conceptual themes that matched interview protocols. Additional codes were developed and applied via content analysis. As an example, one descriptive code (“Decisions Post-PhD”) retrieved every section of an interview where a participant prospectively discussed their future career decision-making and planning.

Our research team case-studied this population utilizing a combination of coding reports and full-transcript critical reads for each participant. Each participant’s case study covered a thorough history of their experiences from pre-PhD years up through PhD, postdoc, and job searching and attainment.

Individual research team members independently took on the initial case study research and write-up. They then presented their case studies to the team, where they were discussed and scrutinized for any additional details that needed to be expanded upon or added. This process of case study creation, group discussion, and case study refinement resulted in thorough case studies verified from many researcher perspectives. These case studies provided the material for the research team to conduct secondary comparative analysis across individuals in the sample.

Using Excel, Researchers (CMH, RFJ, RM, and RR) created a thematic database of all 40 participants in order to assess variance and commonality of themes. Derived from initial case studies and modified as subsequent case studies presented additional themes, the database contained every theme that pertained to scientific development, environment and relationships, and career differentiation. All researchers populated the database for their individual case studies. Two researchers (CMH and RFJ) then expanded themes through analysis and discussion of additional case studies. This iterative approach allowed new themes to be added in several waves. Tertiary analysis allowed themes to be retired when they proved non-relevant to the research aims. Thematic saturation resulted in 42 themes related to career development and outcomes, grouped into 7 categories (Table 2). Themes contained dichotomized or scaled responses (e.g., “yes” or “no”, and “positive,” “mixed,” or “negative”) as well as open-text fields where a summary of that theme as it applied to each case could be written. We tracked most themes longitudinally with fields for PhD and postdoctoral phases.

**Table 2.** Preliminary Themes.

| Category | Themes |
| --- | --- |
| Research Progression | Pre-PhD Science Experiences<br>Level of Independence<br>Progress toward a Scientific Niche<br>Evidence of High Impact Publications/Project<br>Evidence of Significant Project Challenges<br>Published During PhD/Postdoc<br>Obtained Competitive Grants/Fellowships |
| Work Environment & PI/Mentor Relationships | Quality of Work Environment<br>Structure of Lab Promotes/Inhibits Independence<br>Frequency of Contact with PI<br>PI's Engagement with Career Development<br>PI Relationship Quality<br>Has Non-PI Mentors<br>Shares an Identity with Mentors (Race/Ethnicity, Gender) |
| Science Identity, Traits, Behaviors & Roles | Earliest Identification with Science or Scientist Traits<br>Possess Mentor/Teacher Identity (mentor/teacher delineated)<br>Takes Risks in Research or Career Decisions<br>Negotiates/Pushes Back<br>Seeks Support/Mentorship<br>Level of Self-Confidence<br>Engagement with Science Community |
| Influence of Social Identity on Scientific Training & Career | Class/Socioeconomic Status<br>Racial/Ethnic Identity<br>Gender Identity<br>Sexual Identity |
| Career Perspectives | Durable Positive View of Academic Careers<br>Expressed Downsides of Non-Academic Careers |
| Career Planning & Skill Development | Engaged in Career Exploration<br>When Did Career Intentions Emerge?<br>Project Management & Supervisory Skills<br>Mentoring & Teaching Skills<br>Collaborative Skills<br>Grant Skills (writing, seeking)<br>Strategic Planning/Decision Making |
| Core Motivations/Drivers toward Academic Career | Teaching/Mentorship<br>Diversify Science/Academia<br>Scientific Freedom or Autonomy<br>Time Balance/Flexibility<br>Personal Connections to Research Topic<br>Translational Contributions<br>Basic Science Contributions<br>Discovery/Learning/Problem-Solving/Idea Generation |

Table displays the full set of analytic themes concerning participants' progression toward RIFC attainment. Distilled data included a fixed option (e.g., high/medium/low) and a short text summary for each participant. Themes were analyzed longitudinally with fixed options reflecting multiple time points, most often PhD and postdoctoral stages.

Analysis of preliminary themes was an iterative process designed to identify converging primary themes and those which varied. Analyses included compiling counts of fixed choice responses and testing hypotheses about relationships between variables – for instance, between quality of mentorship and likelihood of seeking out mentors beyond the PI or advisor, or relationships among primary drivers to pursue an academic career. The results describe developmental processes, primary drivers, and relationships among factors influencing attainment of RIFCs.

## 3. Results

### Pre-PhD Experiences with Science and Research

The development of interest in science is an unsurprising first step towards choosing to apply and enroll in a biomedical PhD and ultimately the start of any RIFC trajectory. All participants had exposure to science prior to starting PhD training. Those who attained RIFCs in our population, however, had varied pathways to science, research, and ultimately biomedical PhD programs. These pathways were often mediated through familial proximity to science (and socio-economic status). Fourteen participants had parents and/or other close family with advanced degrees and careers in science, medicine, or engineering, a proportion well above the national rate of STEM workers with advanced degrees (12). Of the 26 participants who did not have family working in science or engineering, nine were first-generation college graduates. Those without early familial exposure to science discovered and navigated science and research by drawing on other key resources.

Those with family members who worked in scientific or medical fields were exposed and acculturated to science early. Their families shared their experiences in science, helped foster scientific ways of thinking, and introduced participants to scientists and scientific environments. Caleb abandoned a business career to pursue biomedical science, attributing this decision to his scientist parents and the interactions he had with scientists throughout his life. He said:

> My Thanksgivings were spent sitting around a table with 10 PIs discussing the philosophy of life, essentially. That way of seeing the world and thinking about things is ingrained in me. I’ve always been somewhat curious and wanted to know how things work and why things work, and even though I was originally pursuing a career in business, I was still struck by that.

Christopher grew up in a household with two parents with terminal degrees in science and medicine. He explained how his mother helped him get his first research job, which led directly to his decision to pursue a PhD:

> …my mom works [there] and knows this guy. She’s the person that ended up getting me this job, which has been [a] really important part of my life so far… It’s something that I don’t think I would have gotten if it wasn’t through my mom… I wouldn’t have gone to graduate school if I didn’t have a good experience working in [that] lab.

Those without familial access to science drew on other resources, including family encouragement, teachers and mentors, science classes, and institutional resources and programming. Ayesha, who described herself as a “first-generation scientist,” received encouragement from her mother, who stressed the importance of pursuing knowledge. Ayesha’s mother encouraged her to pursue her first non-classroom science experience in a clinic which then became a main motivator for pursuing a PhD.

> Someone in my family had [genetic disorder] and I was curious as to why he was sick and what was going on. My mom [said], “Try to find the answer yourself.” And so, I literally just called up the hospital, and I talked to someone who worked on [genetic disorder]. He told me to come talk to him, and I started volunteering in the [clinic].

Like Ayesha, others discussed the importance of familial support in their early educational and scientific development. Their families provided encouragement, support with schoolwork, and emphasized the importance of education.

Several participants discussed experiences in school that provided entry into science. Branden, a first-generation college graduate from a low-income household, found mentorship through high school teachers, who helped form his interests in science and teaching.

> …some of my best teachers were science teachers. Just the background of the way I grew up, teachers were the only people I could look up to anyway. And so, they were usually my heroes, and one reason why I’m in graduate school now is that I really want to be an educator.

Other participants similarly shared stories of instructors who cultivated their interest in education and science by serving as role models and providing scientific opportunities.

Others became interested in science through high school coursework. Marshall, who grew up in a lower middle-class family, explained that classroom experiences shaped his scientific aspirations:

> At some point in high school, I just knew that I was gonna be doing something scientific… it just seemed like the only option, because it interested me the most… then when I went into college, I chose to major in biochemistry… I probably chose it based on the courses that were required, and it was a set of courses that I was interested in taking, so it worked out really well for me.

Participants without family members in science often developed their interests by engaging with institutional resources and programming. Cassidy explained how a sequence of opportunities led her to pursue biomedical training:

> My first research experience began as a sophomore in college. My college advisor was doing some work…he asked if I wanted to join him… I learned some basic research techniques and got a little bit of experience… I was in this local biological honors society chapter, and they sent me to a conference to present my research. And while I was there, I just really was impressed with how I enjoyed what I was doing and explaining it and talking to people about science… I applied for a research experience for undergraduate fellowship program…That really gave me a desire to possibly pursue further education in the field. I had never intended upon having any sort of biology career, and that really just made me decide that it would probably be worth pursuing…

Institutional resources and programming provided exposure to science that participants without family members in science might not have received. Moreover, these germinal interactions with science shaped perspectives about what science is for and what scientists can do. The development of interest in a field also serves as a first stage of development of a career perspective.

### Evolution of Career Intentions, and the Perspectives and Motivations behind them

We tracked the evolution of trainees’ career perspectives, motivations, and intentions. As trainees progress, their experiences and exposure lead to refined perspectives and an increased understanding of how their motivations could align with a RIFC. This refinement informed the articulation of their career intentions each year. Participants’ career intentions (Figure 1) followed three patterns: a stable, consistent intention to pursue a RIFC (n=21); an increasing intention to pursue a RIFC (n=12); and fluctuating intention where a goal of pursuing a RIFC was temporarily discarded before returning to it (n=7).

**Fig. 1.**
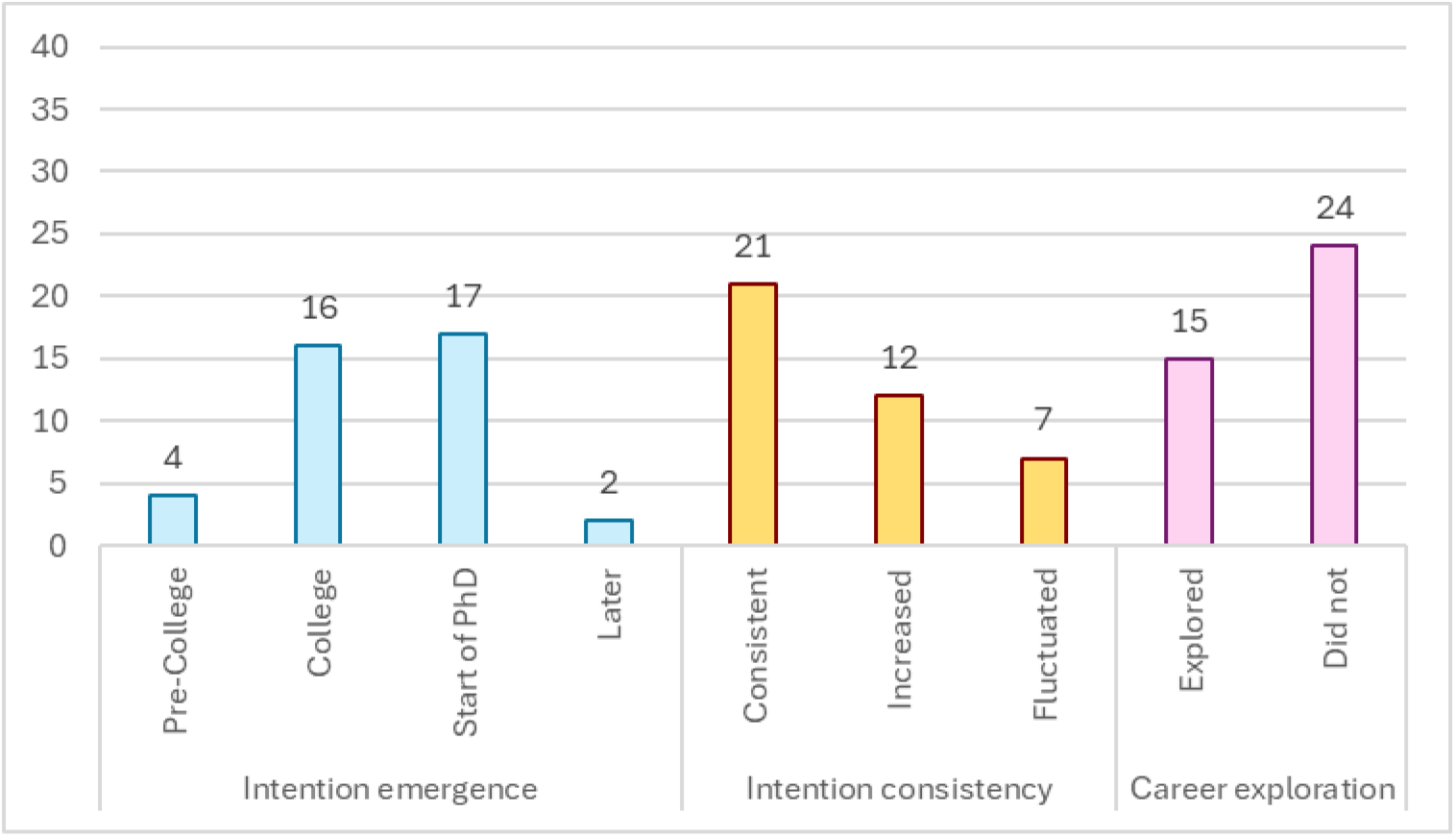
Trajectory of Academic Career Intentions. This figure reports counts for three dimensions of academic career intentions: when intentions emerged, whether intentions were stable or followed a pattern of change, and whether participants engaged in meaningful exploration of other potential careers.

### Evolving Perspectives on RIFCs

Participants frequently detailed their perspectives on various careers. We define perspectives as participants’ holistic views of a career, negative and positive aspects of that career, and how they expect that career will fit into their life. These perspectives shape career planning and decision making. All participants eventually arrived at a positive view of academia, with most participants (n=34) holding an overall positive view throughout their training. As we displayed with early science experiences, perspectives are influenced by exposure to work, those who do it, and relationships with those people. In the case of RIFCs, this means PIs, labmates, and biomedical trainee peers alongside firsthand experiences all shape an individual’s RIFC perspective. Positive perspectives offset and outweighed drawbacks of the career. As Owen explained:

> I think [attaining a RIFC is] really hard; very, very hard, but not impossible. It’s just a lot of work. But I think if you love the work then it’s not gonna be a problem.

A small number (n=6) viewed academia negatively for a prolonged period. These perspectives were typically rooted in structural factors such as pay, job instability, and work/life imbalance as opposed to a dislike of research. Michelle, who considered various careers, expressed serious concerns about the challenges of starting a career as a PI.

> I know of a few friends who are now assistant professors, and it was extremely difficult, especially in the first year, to balance setting up a lab, doing teaching, trying to get projects going, submitting grants. It’s just extremely overwhelming and you’re constantly worried about money.

Over several interviews, Michelle’s appraisal of academia wavered. Despite her reservations, her RIFC intention began to solidify as she gained research success and a supportive research community, defined a niche, and began to focus more on the societal contributions she could make in research-intensive roles.

Even those with consistently positive views of academic careers acknowledged potential drawbacks. Some participants discussed personal circumstances of familial economic support that mitigated the impacts of drawbacks. Logan reflected on the conflict between salary and time demands in academia and his desire to start a family. Yet factors in Logan’s life protected him from these challenges:

> I come from an upper middle-class background and so does my wife, and I think if you were coming from a lower-class background or middle-class background [and] you had a sudden health expense…A lot of people, they may just have to say, ‘Look, I can’t afford this…” All these things can have this accumulating influence on the quality of life and stress for scientific trainees. So, I think that’s where if you come from money and you have that safety net, you can withstand that…

In addition to familial wealth, Logan’s wife’s career had scheduling flexibility that allowed her to assume the bulk of childcare labor. Similarly to Logan, Bruce acknowledged the salary limitations of academia but noted this was not a concern for him specifically because his wife’s salary could support their family. Logan and Bruce exemplify how social-structural factors can impact one’s perspectives and ability to pursue an academic career.

### Drivers Towards RIFCs

Participants described what motivated them towards a RIFC as well as other careers each year. We define motivations as the personal and professional values and desires of a participant as applied to the perceived possibilities of a career. In that regard, a motivation is contingent upon an individual’s career perspective, or what they believe is possible within RIFCs. As their perspectives on RIFCs broaden and become nuanced, an individual has more avenues where they can imagine enacting their values and desires. For example, trainees with little exposure to teaching might form a perception that they cannot fulfill their desire to teach in RIFCs but later find that RIFC mentorship responsibilities fulfill the same passion. If those motivations find footing in a specific career perspective, they act as a core driver towards that career goal.

In this section, we discuss participants’ primary motivations for pursuing a RIFC. Most participants described multiple, high priority drivers, with only one participant indicating just one driver. Across the 40 participants, we identified seven common drivers. The most common drivers were scientific freedom and autonomy (n=28), discovery and idea generation (n=27), contributing to scientific knowledge and human health (n=23), and teaching and mentoring (n=21). Participants were split on whether it was possible to obtain adequate work-life balance and a flexible schedule, but those who saw flexibility as a feature of academic careers valued it as a primary driver (n=14). Participants who rated low for time balance and flexibility as a driver (n=17) identified a lack of balance as a potential drawback to an academic career or were simply not concerned with excessive time demands. A small number of participants (n=7), all women and/or members of underrepresented racial/ethnic groups, were highly motivated to diversify science and saw this work occurring in concert with their roles as a scientist or teacher and mentor (Figure 2).

**Figure 2.**
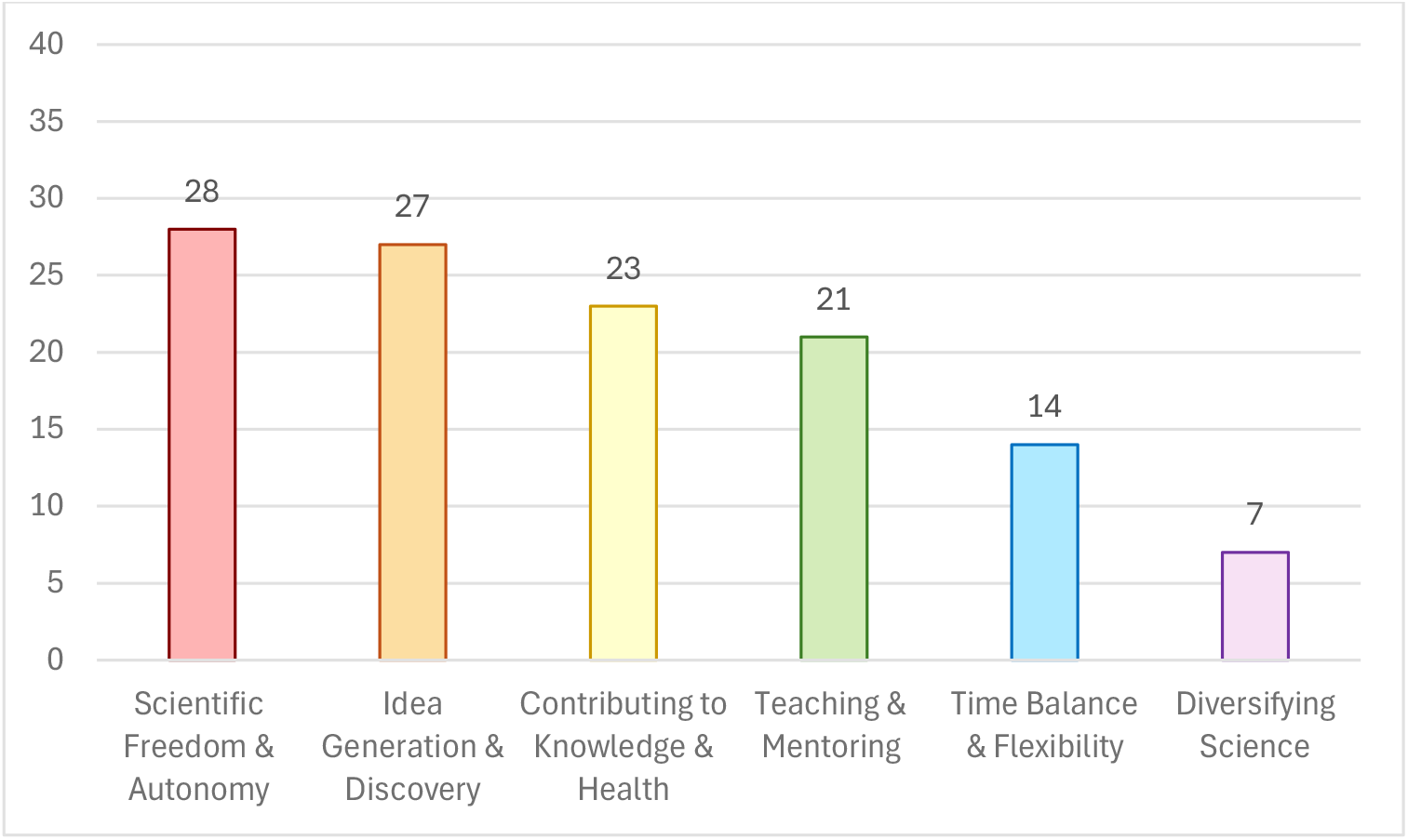
Drivers Towards RIFCs. The figure displays the frequency of drivers towards RIFCs across the population.

### Scientific Freedom and Autonomy

One of the most common drivers (n=28) towards a RIFC is the freedom and autonomy afforded by being an independent investigator. Comments about the importance of autonomy appeared throughout interviews. Bruce explained:

The most appealing aspect of [an academic career] is the autonomy that you have. You get to be the boss and decide where you want the projects to go. If you can convince someone to pay you for it, you can do whatever you want. I think that is the draw for me.

Kevin echoed Bruce’s feelings and compared the freedom of academia to a career in industry.

> The most attractive thing [about] academia is the freedom you have to conduct any kind of research you want. The freedom is the most attractive thing. If you go to industry, it’s always project oriented. You cannot choose what you want [to do]. If you’re not at a manager level, you cannot handle things in your [own] way.

Scientific freedom and autonomy were generally discussed in two ways: having the intellectual freedom to determine one’s own projects and research direction and developing one’s own work style without direct supervision.

### Idea Generation and Discovery

The ability to generate ideas and regularly seek novel findings was inherently motivating for more than half of participants. They often described lifelong, innate curiosity and a passion for uncovering and enhancing scientific knowledge. These participants were drawn to the process of scientific discovery, even when this process was not linked to a specific application or outcome. Lori stated:

> I really like science and discovery, so I want the freedom to really have good ideas and be able to follow through with them… A successful career would be one where I could do research on things I thought were interesting and important… not [finding] a treatment tomorrow.

These participants valued the scientific process and the experience of discovering new knowledge apart from the impacts of their science. Timothy, who initially sought a teaching-intensive career, described answering original questions as a primary factor strengthening his intention to become a PI. He said:

> More and more, I’m leaning towards being a principal investigator at an academic institution. I like my job. I like thinking about experiments. I like coming to work and doing those experiments. I like analyzing data and thinking of questions and thinking about how to answer those questions in a way that leaves no doubt in your mind.

For well over half of participants, a fundamental draw of a RIFC is the ability to generate questions and ideas leading to new knowledge. This specific goal is separate from seeing the impacts of one’s science.

### Contributing to Scientific Knowledge and Human Health

Twenty-three reported that a primary driver for pursuing a RIFC was making meaningful contributions to scientific knowledge and human health. Timothy articulated his decision to pursue a PhD over an MD in the ability for his basic research to help people downstream.

> As an MD, you might be able to treat thousands of people during [your] career. But as a basic biomedical researcher, you’ll be able to help millions of people throughout your career, so that’s how I rationalized [my decision].

Virginia commented on a draw of a scientific career:

> …Creating knowledge is amazing, helping other people… if I can do anything to try to prevent cancer patients from dying…

While many participants spoke of a love of discovery and generating knowledge, those who sought to contribute to scientific advances emphasized the ability to help people via knowledge production. Most participants aligned with this driver spoke about the potential impacts of their research, even if they were not directly working on therapeutic applications or interacting with patients in clinical settings. Among participants in this population, the desire to have their discoveries contribute to human health overrode the distinction between basic and translational research – both types of researchers are motivated to improve the lives of fellow humans, no matter how proximate or distant.

### Teaching and Mentoring

More than half (n=21) indicated that a major draw of a RIFC is the opportunity to teach and/or mentor, with some indicating early that it was their main objective in pursuing a career in academia. As these trainees progressed through their PhDs and postdocs, those who initially focused on teaching or mentoring were flexible in refocusing their aspirations to suit different teaching and mentoring settings. For example, two participants initially interested in classroom teaching saw themselves fulfilling a similar pedagogical role by teaching and mentoring in the laboratory. Some saw teaching and mentoring as a method of supporting students and/or promoting diversity in science.

Branden began his PhD career with a strong interest in teaching at a liberal arts college. He referenced his love of teaching throughout his PhD and postdoc. Early in his PhD, however, he recognized the potential for research to be used to teach and mentor outside of the classroom. Branden commented on this philosophy:

> I’m interested in research as a pedagogical tool… to teach [students] what science is all about and exactly [why it] is worth doing as opposed to more classroom-oriented sorts of things.

Later in his PhD, Branden described how his successes in graduate school opened doors for him to become a PI at a research-intensive university, and how his goal shifted away from teaching at a liberal arts college. At the beginning of his faculty career, he viewed advising graduate students on professional development as a part of his educational work, even using “teach” to describe advising and mentoring activities.

> What I really think [is] the point of graduate education…is the critical thinking stuff [and] the professional development… That’s what it is that I’m trying to accomplish when I’m working with [graduate students].

Janette identified an interest in classroom teaching early on, but as her career goals shifted towards academic research, she discovered she enjoyed mentoring in the lab and found the role to be more compatible with that of a PI. During her postdoc, she said:

> I really am enjoying mentoring right now. I took on a graduate student. I’ve really kind of put her under my wing. We discuss different things she’s doing and [I’m] her go-to person, and that’s been fun, just to feel like I’m almost a PI in that sense…I’m finding that as rewarding as I did some of my previous teaching that I had done.

Though teaching and mentoring are distinct fields of practice, each with its own philosophies and associated competencies, biomedical scientists do not always draw bright lines between them. Many viewed their desire to engage with students as transferable to different mentoring and teaching contexts.

### Time Balance and Flexibility

Participants were split on their interpretations of time demands in academic careers. Fourteen perceived the career to be flexible, with the ability to control their time to fulfill other obligations, despite the expectation of long hours. In contrast, 17 felt that time balance and flexibility were disincentives, due to long hours and an inability to have work-life balance.

Robert saw the flexibility of an academic career as a primary motivator and found himself more scientifically productive when he had the flexibility to control his work time and balance his work with family obligations.

> [Directing] my work is a big [motivator], but I think the biggest thing for me is, because I have a family, having flexibility in my work life is incredibly important…I’m far more productive when I do have more flexibility.

The pairing of scheduling flexibility with a self-directed work structure was common among the group that highlighted flexibility. All but one of the 14 who viewed schedule flexibility as a draw were also highly motivated by scientific freedom and autonomy, often uniting these drivers under a desire to “be your own boss.”

In contrast to participants like Robert, 17 did not cite balance and flexibility as a draw to RIFCs and were instead motivated by other factors. This group tended to focus more on work-life balance than flexibility, with several participants discussing the downsides of long hours and limited time for personal goals and responsibilities. For Denise, academia compared unfavorably to corporate environments regarding work-life balance:

> It’s not great, the career life…I would like to have more time to spend with my family and the people I love…I can definitely see people that work in the private sector, you know, they have set hours, they work eight to five and then they go home, sometimes I go home and I still have work to do…[It’s something] I fear because…in academia it’s so competitive…If I follow the path that I want which is to become a faculty member, it’s going to be difficult…

Denise’s focus on balancing work with family was not uncommon among others who viewed the time demands of academia as a drawback. In addition, some highlighted concerns about managing existing health issues or preventing the negative consequences of long work hours. Notably, there were no gender differences among those who valued flexibility, nor among those who identified a lack of flexibility as a drawback.

A small number were more explicit that long hours would not pose a challenge. They were less focused on work-life balance or time flexibility as motivators or detractors. Branden stated:

> I probably work about 50 hours a week, I would guess, in some variety or another. Some of that might be at home doing work on the computer. I mean, they are long hours. They’re not so long that I am feeling burned out or anything… I think the amount I’m working is probably about what’s expected, and so I don’t really have any complaints about that.

Despite participants like Branden, who are less concerned with the time demands of RIFCs, our findings reveal a split between those who value flexibility and those who value work-life balance. The close pairing of time flexibility and scientific freedom as motivators toward RIFCs suggests that some trainees view time considerations through a general desire for autonomy. Others saw time considerations as a balancing of demands between their desired career and their personal lives and non-work aspirations.

### Diversifying Science

Diversifying science was a primary driver for seven participants. These participants felt they could make a difference and promote belonging and persistence among historically underrepresented students. Those drawn to diversifying science were women and/or members of historically underrepresented racial and ethnic groups. Darian viewed diversifying science as a social responsibility, explaining:

> A big part of my science career [is] aggressively making sure that I can be a part of a change to make sure that people are more aware of the options in science for African American men and women.

Participants employed a variety of strategies to support underrepresented students. Some, like Melinda, acknowledged the dearth of professors from their racial and ethnic groups during their own education and training, and how their presence in academia can signal the possibility of succeeding in the professoriate.

> Going into academia means a lot more for me than if I wasn’t Hispanic… I think less than ten percent of tenured professors are Hispanic. It’s crazy, considering the population of the US… I need to make this happen, because then I can have more of an impact on undergraduate students and retaining grad students from Spanish backgrounds.

These participants were active and creative in their efforts to promote diversification. One participant chose a position where she grew up in order to offer scientific training to her community. Another had experience with and planned to continue developing initiatives to promote science within marginalized communities. Several participants even published work on racial/gender dynamics in PhD and postdoctoral training.

A few participants noted their efforts to promote diversity were not always rewarded or supported. Samantha described an experience where her PhD PI felt her diversity and outreach work took time away from her science. She explained that for her, and for other Black scientists, “giving back” to community is a core element of her identity and her reasons for pursuing a RIFC:

> [My PI] feels like I had committed more towards diversity and outreach work than I had towards my science, and that was very difficult for me to hear. This has been a very common theme, at least amongst my associates who are from [a] Black background or even URM [underrepresented minority] backgrounds who are in science, that a lot of times it’s viewed as if we are spending more time on diversity or community outreach work than we are on science. Not knowing that a lot of us, we just double our hours and we do both…[Part of] our identity as an educator or as a scientist or as a scholar is giving back…It turned into a situation where I had to kind of end the conversation and get up and walk out of the office because I needed to digest what he had just told me.

Denise also faced challenges with her goal to support underrepresented students. She started a PhD with the hope of returning to her home country to teach and mentor students. During training, economic shifts in her home country led her to believe that a professorship was no longer an option there. In response, she discontinued her efforts toward an academic career. Through a chance event, she volunteered in mentorship and teaching roles in her home country, which restored her passion for academia. She explained:

I want to be a part of the network here in [country of origin]. Because I think here’s where I can make the impact with the students…I work voluntarily from Monday to Wednesday in the lab of my undergraduate mentor and then Thursday and Friday I teach…When I’m here, I see that I love working with the students…it was like reconnecting with that first love that I had. That was the whole reason why I went to grad school. Somewhere in the journey I got discouraged about the system and I didn’t see myself as being a faculty member in the States… Then that left me with, ‘Okay, now what am I going to do with my PhD?’

Denise’s story illustrates how academic career plans that include a focus on diversifying science can be fragile when opportunities to engage with diversity work are restricted. As Samantha noted above, those with a dual focus on research and diversification encounter challenges like increased workloads and a lack of appreciation for their efforts. Together, Denise and Samantha’s accounts emphasize the importance of supporting trainees who are motivated to work towards diversification so they can continue to pursue academic careers.

The seven who placed a high priority on diversity work had other high priority drivers, often teaching and mentoring and scientific freedom and autonomy. Yet their commitments to diversifying the scientific workforce should not be understated. Rather than view diversity work as being service-related or superfluous to their academic pursuits, these participants viewed this work as a primary motivator for pursuing academia.

Nearly all participants were drawn to RIFCs for more than one reason. Many themes appeared with high frequency, and some drivers frequently appeared together. For example, many expressed drivers in combination, including scientific freedom and idea generation and discovery (n=20), scientific freedom and time flexibility (n=13), and teaching/mentorship and idea generation and discovery (n=14). However, developmental themes and primary drivers come in many combinations, with most participants expressing three high-priority drivers. Participants’ primary motivators sometimes evolved over time, such as shifting from teaching to research. Participants were more consistent with other drivers, like freedom and autonomy. In this population, what drives participants to a PI career is a shifting set of constellations drawn from a set of common themes.

#### Career Intentions

Career perspectives offer the scaffolding for motivations to attach, and together they shape career intentions at every interview. We define career intentions as a combination of career interest and planning. Twenty-one described a steady, consistent intention to pursue a RIFC (Figure 1). They settled on this intention early and did not meaningfully waver. Occasionally, participants in this group discussed other careers and compared academia favorably to them. Ayesha said:

> I’m just not an industry type person. I’ve tried to explore types of things like this because it would keep my options open. And I just feel like there’s a part of the critical thinking or the creating that [industry is] missing. That’s why I really like [academic] research. Even though [there is] a lot of failure, it’s satisfying failure.

Others had little to no interest in considering other career paths or actively avoided career exploration to maximize their focus on research-intensive roles. When asked whether he had a “backup” career in mind, Jeffrey explained:

> Let’s say someone wants to be a science writer. They’re writing blogs on something that might help them with journalism in the future and picturing that kind of career. I think if you’re excited about academia, you don’t have time to do that stuff. It’s not gonna contribute to your academic career.

In contrast, about half of the participants changed their career intentions during training. A quarter had fluctuating intentions to pursue a RIFC, often decreasing during a period of career exploration before becoming their primary intention once again. For example, some participants with fluctuating intentions started graduate school with a RIFC intention but later discovered unappealing aspects of RIFCs, such as the degree of competition and grant writing, and chose to explore other careers. Forces such as external validation or exposure to positive aspects of RIFCs led them back to their initial RIFC intention.

Twelve of the 40 started training without a primary intention in a RIFC but gradually increased their interest and achieved that career. Those whose interest increased often began with a focus on teaching-intensive faculty positions or industry, and later realized that their research interests, successes, and strengths better aligned with a RIFC. The process of increasing interest was varied and influenced by several factors, but participants often described how their growing attraction to aspects of RIFCs informed their evolving intentions. For the first several years of training, Laurie viewed academia negatively, believing its demands would prevent her from being able to enjoy research:

> I really don’t want to have my own lab… I like doing the research, but I feel like I don’t really want to be in charge of everyone else’s research, in charge of writing grants and worrying about funding, and I feel like I would just be way too stressed out, and wouldn’t have enough time to actually do what I like doing in the first place…

In subsequent interviews, Laurie described her late-stage switch towards a RIFC as the result of several factors, including external validation and encouragement, thorough exploration, and shifts in priorities as well as her relative perceptions of academia and industry. She was encouraged to remain open to academia:

> …my boss still thinks there’s hope that I’ll change my mind… It’s more or less, “Oh, you never know… you might change your mind if the funding gets better,” …people are still pushing for people to stay and pushing for me to stay. And also, family friends that are both PIs, and they said some more things… that “it’s not that bad, you’ll like it.”

Another participant, Janette, began graduate school with a strong desire to teach at a liberal arts college, believing that this career would offer the best balance with her goal of starting a family. During her postdoc, she began to perceive tension between teaching careers and family life, a shift in her perspective which led her to focus on preparing for a RIFC instead.

> …my first year, I said I wanted to teach. That’s slowly become more of a non-reality, or interest, especially if I want to be able to maintain a family life and a career. Teaching is -- those that can do it well, lose some other balance in their life, and family life is not something that I’m willing to compromise on.

Janette’s changing perspectives on teaching roles led her to reconsider her perspectives on work-life balance in RIFCs.

Not all participants in this population began their training journeys with a positive view of academia or a strong intention to pursue a RIFC. About half of participants’ career intentions changed over time, with a significant number increasing or regaining interest in RIFCs after having ruled out this career.

#### Mentoring and Its Influence on Career Differentiation

While examples of PI relationships are interspersed as examples of perspectival shifts, PIs play varied and vital roles in the process of career differentiation for our RIFC population. Vicariously, they serve as models for what the career entails and how to balance it with non-work life. Directly, they offer an environment for training, skill building, and external validation.

Those with positive mentoring relationships described myriad qualities tailored to their individual situations and work styles. These qualities included accessibility, a sustained interest in their work, constructive feedback, being a role model professionally and socially, career development support, and research and career opportunities. Casey described his two postdoc mentors as supportive of career development and attributed his success as a PI to their guidance.

> [My advisors are] quite complementary to each other…Both of those people really brought different perspectives to how to do science. Also, they just had a lot of experience in terms of, how do you write grants, how do you give a job talk, how do you interact with people at meetings, and I’m still learning all these things. I don’t think I would be where I am in my career without having that exposure.

Some participants had a PI who was too busy to provide support. Nikki’s PhD PI was often unavailable, which slowed her progress.

> She goes out of town for conferences, and she recruits because she’s a program director and so she’s not always there for me to ask [questions]. That kind of gets frustrating sometimes…I shouldn’t complain about it so much, but I [have] had to look up a lot more things on my own.

Strikingly, 30 participants were proactive about seeking additional mentors, regardless of the quality of their primary mentoring relationships. Melinda’s postdoc PI was neither available nor supportive of her scientific and professional development. To compensate for inadequate mentoring, guidance, and encouragement, she sought and relied on others.

> I found a mentor in a different department, and she’s been such a source of everything I’ve dreamed about and more from a postdoc mentor…she’s been so supportive, helping me in so many ways, helping me with my job interviews, helping with my negotiations, everything…You kind of get jaded and forget there are really great mentors out there.

In addition to her new mentor in a different department, Melinda connected with mentors outside of her “cutthroat” institution, which allowed her to see a larger, supportive scientific community.

Acquiring experience to not only carry out science but to understand what the role of a PI can entail is a gradual process. Our analysis reveals that these developmental processes begin from the earliest exposure to science and continue through the postdoc and faculty transition. PIs and mentors play a critical role not only as supporters in scientific development but also as models for what RIFCs can look like. Those bound for RIFCs share common developmental experiences, including pre-PhD research experiences and proactive mentorship seeking.

## Discussion

By focusing on developmental processes, we identified key aspects of career differentiation— perspective formation, motivation fit, and career intentions. Career perspectives develop over the years of a PhD and postdoc through exposure and experience. When perspectives align with an individual’s values and desires, they motivate them towards that career. Those motivations, in turn, are vital in creating career intentions. For this population, we found that fluidity of career intentions was common and for some persisted into the postdoctoral years. Additionally, we identified six common career motivations that varied in their combinations. Early exposure to science was mediated by class and by having family employed in scientific careers; those without familial proximity to science used a variety of strategies to obtain science experiences. Mentors often played important roles in shaping perspectives and offering support for science and professional development.

Our data show the importance of early exposure to science and research for generating interest biomedical training and careers while also forming the germinal perspectives of holding a scientific career. While nearly all participants completed pre-PhD research experiences, their earliest experiences with science took many forms. Fourteen participants (35%) had close family members with advanced degrees working in science, engineering, and medicine, a rate that exceeds the 20% of the national STEM workforce holding advanced degrees (12). These participants’ families played crucial roles in their pathways into science by providing exposure, fostering scientific traits, and facilitating professional connections. Given that participants used firsthand experiences and observations of role models to form RIFC perspective, this exposure may have additionally provided more nuanced scientific career perspectives at a younger age.

A higher fraction did not have parents in the STEM workforce and/or were the first in their families to receive a college degree. These participants proactively sought a range of entrées into science. Encouragement from family and supportive teachers was a vital component of gaining both exposure and experiences in science and research. Literature shows that receiving encouragement to pursue science by family and teachers predicts formation of scientific career aspirations (13) and makes science knowable, “thinkable,” and aspirational (14). Outreach efforts could extend to families and high school teachers to raise awareness of biomedical career paths.

Participants also utilized institutional programming and resources, such as engaging science curricula and research opportunities, to gain entry into science. Our findings suggest the importance of scientific outreach and accessible research experiences (15–16). While there is much research covering individual supports and points of exposure (17), our data point to multiple formal and informal pathways into science that must be considered holistically.

### Many participants started their training with no intention of seeking a RIFC, or changed their career intentions several times before deciding on a RIFC

By utilizing a longitudinal design, we captured participants’ shifting perspectives on academic careers and the stability or change in their career intentions over many years. Career intentions shifted for about half of participants over the course of their PhD training and postdoctoral years, with a significant proportion of participants increasing their interest toward RIFCs. For many whose interest in RIFCs increased, the shift involved an initial interest in another career with teaching-intensive faculty being the most often desired career. About a quarter explored and considered other careers at some point between the start of their PhD and the start of their faculty career. While singularly PI-goal-minded trainees are easier for mentors to identify as faculty-bound, evidence of those who increased from and explored other careers points to the prevalence of non-linear paths toward faculty careers. These shifts happened at all time points which further underscores that there is no singular timepoint predictive of what career someone will set as a goal or attain. Many shifts began or continued into postdoc years, suggesting that the postdoc stage has a distinct key role in the development and crystallization of RIFC perspectives and intentions. Supportive mentors should embrace this reality and remain receptive and responsive to movement both toward and away from RIFCs. Mentors should provide opportunities for all trainees to develop the skills needed for a RIFC. Trainees should seek out and be exposed to many models of what a RIFC can look like by those who hold the position, further grounding career perspectives, allowing for a sense of fit.

### Participants described common motivations for pursuing RIFCs, but in different combinations

Research on the factors influencing persistence towards a RIFC in biomedicine is underdeveloped. Through a grounded reading of each participant’s interviews, we identified seven main drivers propelling participants toward RIFCs. In a landscape in which many biomedical trainees elect not to pursue RIFCs, identifying motivations toward RIFCs and the components of RIFCs that those motivations can attach to is key for understanding what influences persistence.

In line with the literature (18), “scientific freedom and autonomy” and “discovery and idea generation” were the most common drivers. These are both commonly cited as aspects of a scientific career, even by those with minimal proximity to them. These consequently may be the most common drivers that mark initial interest and then endure throughout training and career attainment. However, “teaching and mentoring” appeared nearly as frequently. A key discovery in our research is the enthusiastic and flexible attitudes towards teaching and mentoring among a population ultimately destined for research-intensive roles. About half of the 40 began their training journeys with a strong attraction to teaching and mentoring, maintaining it throughout. While their intentions shifted gradually toward RIFCs based on their enjoyment of the research process and/or their research success, some reframed their desire to teach as a desire to mentor and train others. Other individuals did not draw strong lines between mentoring and teaching, focusing more on supporting students. This tendency to view teaching and mentoring as malleable, and to embrace the idea that mentoring and training can fulfill various supportive and pedagogical goals, has not been demonstrated in the literature.

Participants identified other strong drivers, such as a desire to diversify science, common among women and people of color. Participants’ enthusiasm for teaching, mentoring, and diversifying science suggests that many aspiring PIs hold commitments beyond research and scientific discovery. While non-research passions were common, participants sometimes experienced friction between these passions and the traditional expectations of training. Welcoming dialogue about the many passions that drive RIFC aspirants would serve broader goals of the scientific community such as diversification of the professoriate and cultivation of the next generation of scientists. Additionally, continued research on early-career faculty to see how their drivers align with workplace values and rewards is needed.

Another novel contribution of our research is parsing the desire for time flexibility from the desire for work-life balance. Both women and men discussed the challenges of balancing career with family and the social supports required for sustaining a career while having a family. We saw variation in our population in the extent of social support for maintaining work-life balance, including familial help with child rearing and financial support. These findings echo the importance of providing support for maintaining work-life balance among biomedical trainees (11).

However, a popular driver among this population was the flexibility afforded by an academic career. Many commented on how the structure of a RIFC allowed them to choose their own hours and dictate their time, leaving them more flexibility for structuring their non-work time. Delineating flexibility from balance in discussions of RIFCs would support trainees in making informed decisions about which careers align best with their responsibilities and lifestyles.

Our research on drivers and motivations uncovers what allows trainees to endure a highly uncertain career path. They persist based on core beliefs of what value a scientific career can bring to society, the scientific enterprise, and their own fulfillment. These diverse drivers and motivations appear in different constellations for every individual. They are the beliefs, values, and desires of each participant working to find a home in a chosen career. Mentors should engage in candid discussions with trainees about what motivates and inspires them, and what careers would be most efficacious and fulfilling.

### Limitations and Future Directions

One limitation of centering the analysis on those who attained RIFCs is that we lack a comparison group of individuals who pursued but did not achieve this career outcome. That said, the broader study population includes 52 participants who expressed intentions to pursue a research-intensive faculty career at graduation but did not ultimately secure one. While key components of career differentiation were explored in this paper, it will be revelatory to find what factors in the process vary for those who don’t go on to pursue or attain RIFCs. For example, we can ask questions such as: is this outcome a result of desiring but not attaining a RIFC, or do participants in this comparison group continue to differentiate toward non-RIFC careers? Such a study will also help establish what combinations of experiences, perspectives, motivations, and intentions are unique to this PI population and which are not. This future study will also illuminate those factors that promote persistence and shed more light on the choice points many people face in RIFC-striving.

## Acknowledgements

We extend our thanks to the many researchers who have worked on our team, making this research possible since 2008. Dr. Jill Keller helped initiate the studies and her qualitative expertise shaped the project from the beginning. We are indebted to the invaluable work of Fatimah Bhatti, Bryan Breau, Adriana Brodyn, Lynn Gazley, Toni Gutierrez, Sandy LaBlance, Nicole Langford, Ebony McGee, Elizabeth Morrissey, Michelle Naffziger-Hirsch, Letitia Onyango, Jennifer Richardson, Bhoomi Thakore, Simon Williams, Veronica Womack. We extend an additional thank you to Dr. Jeff Engler for insights and suggestions on an early manuscript draft.

## Author Contributions

Conceptualization: R.F.J., C.M.H., C.V.W., P.B.C., R.M.

Methodology: R.F.J., C.M.H., R.R.

Software: R.F.J., C.M.H.

Validation: R.F.J., C.M.H., C.V.W.

Formal analysis: R.F.J., C.M.H., C.V.W., R.R., D.P.O’N., J.F.M., Y.K.C., R.M.

Investigation: R.F.J., C.V.W., R.R., P.B.C., A.E.S., R.M.

Data Curation: R.F.J., C.M.H.

Writing - Original Draft: R.F.J., C.M.H., C.V.W., R.M.

Writing - Review & Editing: R.F.J., C.M.H., C.V.W., R.R., D.P.O’N., P.B.C., J.F.M., Y.K.C., A.E.S., R.M.

Visualization: R.F.J., C.M.H.

Supervision: R.M.

Project administration: R.M., R.F.J., C.M.H., C.V.W., A.E.S.

Funding acquisition: R.M., P.B.C.

